# Loss of RNase J leads to multi-drug tolerance and accumulation of highly structured mRNA fragments in *Mycobacterium tuberculosis*

**DOI:** 10.1101/2022.02.13.480260

**Authors:** Maria Carla Martini, Nathan D. Hicks, Junpei Xiao, Thibault Barbier, Jaimie Sixsmith, Sarah M. Fortune, Scarlet S. Shell

**Affiliations:** Department of Biology and Biotechnology, Worcester Polytechnic Institute, Worcester, MA, USA; Department of Immunology and Infectious Diseases, Harvard T. H. Chan School of Public Health, Boston, MA, USA; Program in Bioinformatics and Computational Biology, Worcester Polytechnic Institute, Worcester, MA, USA

**Keywords:** *Mycobacterium tuberculosis*, RNase J, drug tolerance, drug resistance, RNA degradation, PE/PPE genes.

## Abstract

Despite the existence of well-characterized, canonical mutations that confer high-level drug resistance to *Mycobacterium tuberculosis* (Mtb), there is evidence that drug resistance mechanisms are more complex than simple acquisition of such mutations. Recent studies have shown that Mtb can acquire non-canonical resistance-associated mutations that confer survival advantages in the presence of certain drugs, likely acting as stepping-stones for acquisition of high-level resistance. *Rv2752c*/*rnj*, encoding RNase J, is disproportionately mutated in drug-resistant clinical Mtb isolates. Here we show that deletion of *rnj* confers increased tolerance to lethal concentrations of several drugs. RNAseq revealed that RNase J affects expression of a subset of genes enriched for PE/PPE genes and stable RNAs and is key for proper 23S rRNA maturation. Gene expression differences implicated two sRNAs and *ppe50-ppe51* as important contributors to the drug tolerance phenotype. In addition, we found that in the absence of RNase J, many short RNA fragments accumulate because they are degraded at slower rates. We show that the accumulated transcript fragments are targets of RNase J and are characterized by strong secondary structure and high G+C content, indicating that RNase J has a rate-limiting role in degradation of highly structured RNAs. Taken together, our results demonstrate that RNase J indirectly affects drug tolerance, as well as reveal the endogenous roles of RNase J in mycobacterial RNA metabolism.

## INTRODUCTION

More than a century after the discovery that *M. tuberculosis* (Mtb) is the causative agent of tuberculosis (TB), this disease remains one of the major health challenges worldwide. In 2020, about 10 million people developed TB and 1.3 million died of the disease, positioning TB in the top 10 causes of death worldwide (WHO, 2020). Due to drug penetration issues and the presence of drug-tolerant Mtb populations in human lesions, anti-TB drug regimens must be administered over long durations (Dheda *et al*., 2018, Pontali *et al*., 2018, WHO, 2020). Drug treatment itself may induce further drug tolerance (Goossens *et al*., 2020), and the long treatment period provides opportunities for acquisition of mutations leading to antibiotic resistance. The emergence and spread of multidrug-resistant (MDR) TB are major concerns as it frequently leads to treatment failure and death. In 2020, 7.5% of the new TB cases were either rifampicin-resistant, MDR, or extensively drug resistant (XDR) (WHO 2021) prompting the need to improve TB therapies and prevent the emergence of resistance.

Although the genes encoding the drug targets and activators in mycobacteria are well known, the mechanisms driving the development of high-level drug resistance are less well understood. In addition to high level resistance, Mtb exhibits other forms altered drug susceptibility that allow populations of bacteria to survive for extended periods of time in the presence of antibiotics and apparently serve as reservoirs for the eventual acquisition of high-level drug resistance-conferring mutations (Zhu *et al*., 2018, Safi *et al*., 2019). Initial studies on mycobacterial drug tolerance focused on rare persister cells that are drug-tolerant due to stochastic growth cessation (Keren *et al*., 2011, Kester *et al*., 2014, Harms *et al*., 2016). Further work has demonstrated that altered drug susceptibility in Mtb can arise through multiple mechanisms, which differ in terms of frequency, duration and magnitude of effect. For example, during infection Mtb responds to environmental and metabolic conditions by shifting to slow- or non-growing states in which it is less sensitive to many drugs (Gengenbacher *et al*., 2010, Nandakumar *et al*., 2014, Trivedi *et al*., 2016, Lim *et al*., 2021). Recently, several groups have taken a population genomics approach to identify clinically relevant stepping-stone mutations that facilitate the acquisition of high-level drug resistances. Mechanistic dissection of these mutations has revealed the importance of these different forms of altered drug susceptibility For example, recent studies of Mtb have revealed unexpected forms of genetically-encoded low-level drug resistance such as mutations in *dnaA* and Rv0565c that cause low level isoniazid and ethionamide resistances, respectively (Hicks *et al*., 2019, Hicks *et al*., 2020). Mutations in *prpR*, in contrast, were found to increase tolerance to multiple drugs without affecting resistance (Hicks *et al*., 2018).

We and others have reported genome-wide association studies (GWAS) in cohorts of Mtb clinical isolates (Zhang *et al*., 2013, Hicks *et al*., 2018, Farhat *et al*., 2019, Lai & Ioerger, 2020) and other bacterial pathogens (Diaz Caballero *et al*., 2018, Ma *et al*., 2020, Weber *et al*., 2021) from around the globe. Many of the Mtb studies have identified drug resistance associated mutations in Rv2752c, here called *rnj*, encoding the ribonuclease RNase J. This enzyme has both endonuclease and 5’ to 3’ exonuclease activity and is involved in the 5’ end processing of ribosomal RNAs in *M. smegmatis* (Taverniti *et al*., 2011). RNase J activity is essential for growth in several gram-positive bacteria that do not encode RNase E, in contrast to most gram-negative bacteria which encode RNase E and lack RNase J (Even *et al*., 2005, Redko *et al*., 2013, Cavaiuolo *et al*., 2020). Unusually, mycobacteria encode both RNase E, which is essential and seems to play a rate-limiting step in degradation of most mRNAs (Sassetti *et al*., 2003, Taverniti *et al*., 2011, DeJesus *et al*., 2013, Płociński *et al*., 2019), and RNase J, which is non-essential (Sassetti & Rubin, 2003, Griffin *et al*., 2011, Taverniti *et al*., 2011, DeJesus *et al*., 2017). RNase J has been shown to participate in 23S rRNA processing (Taverniti *et al*., 2011) but we do not understand its impact on bacterial cell physiology and lack a model to mechanistically understand its relationship to drug responses. Here we investigate the role of RNase J in Mtb RNA metabolism and drug sensitivity. We show that *rnj* variants impact drug tolerance and mechanistically link the changes in drug susceptibility to altered gene expression and transcript degradation.

## MATERIALS AND METHODS

### Bacterial strains and growth conditions

*M. tuberculosis* H37Rv and its derivatives were grown in Middlebrook 7H9 broth supplemented with 10% OADC (0.5 g/L oleic acid, 50 g/L bovine serum albumin fraction V, 20 g/L dextrose, 8.5 g/L sodium chloride, and 40 mg/L catalase), 0.2% glycerol and 0.05% Tween 80. Liquid cultures were grown in 50 mL conical polypropylene tubes at 37 °C with a shaker speed of 200 rpm except when indicated otherwise. For growth on solid media, Middlebrook 7H10 supplemented with 0.5% glycerol and OADC was used. For *M. tuberculosis* auxotrophic strain mc^2^6230 (Δ*panCD*, Δ*RD1*, (Sambandamurthy *et al*., 2006)) pantothenate was added to 7H9 or 7H10 to a final concentration of 24 μg/mL. When required for resistant bacteria selection or plasmid maintenance, the following concentrations of antibiotics were used: 25 μg/mL kanamycin (KAN), 50 μg/mL hygromycin (HYG) or 25 μg/mL zeocin (ZEO). Knock-out strains were constructed using the recombineering system described by Murphy and collaborators (Murphy *et al*., 2015). For genetic complementation, an L5-site integration plasmid was used, and for gene overexpression the episomal plasmid pMV762 was used (Steyn *et al*., 2003). A description of all strains used in this study is provided in Table S1.

For growth of Mtb mc^2^6230 in minimal media, log phase cultures grown in minimal media were sub-cultured to an OD600nm=0.01 in the same media. The minimal media had the following composition: 0.5 g/liter asparagine, 1 g/liter KH2PO4, 2.5 g/liter Na2HPO4, 50 mg/liter ferric ammonium citrate, 0.5 g/liter MgSO4·7H2O, 0.5 mg/liter CaCl2, and 0.1 mg/liter ZnSO4. Minimal media was supplemented with 0.05% tween, 24 μg/mL pantothenate, and 0.2% glycerol.

### Antibiotic susceptibility testing

To determine the minimum inhibitory concentration (MIC) for Mtb mc^2^6230 strains the agar proportion method was used (Sirgel *et al*., 2009). Briefly, antibiotics were added to 7H10 plates to obtain the following concentrations: 1, 0.5, 0.25, 0.125, 0.06125, 0.0306, 0.01531 and 0.0077 μg/mL of rifampicin (RIF) or isoniazid (INH). Direct 5 μL aliquots and serial dilutions of 7H9 mid-log cultures of each strain were plated on antibiotic-containing and antibiotic-free plates. The MIC for each strain was determined as the drug concentration that reduced CFU by 90% compared to the control.

### Drug killing experiments

For Mtb mc^2^6230 and H37Rv strains, log phase cultures grown in 7H9 were diluted to an initial OD600nm=0.1 in triplicate in absence of antibiotics. After 24 hours the following antibiotics were added to the indicated final concentrations: 0.6 μg/mL of RIF, 2.4 μg/mL of INH, 2.5 μg/mL of clarithromycin (CLA), 1 μg/mL of ofloxacin (OFX), 2 μg/mL ethambutol (EMB), or 500 μg/mL erythromycin (ERY). Triplicate cultures of each strain were incubated in absence of drug as a control. Aliquots were periodically taken, and serial dilutions plated on 7H10 agar plates without drug. CFUs were counted after 20-35 days.

### Determination of the fraction of survival in INH at different growth phases

Log phase cultures of Mtb mc^2^6230 WT or Δ*rnj* were sub-cultured to an OD600nm=0.02. A total of 24 tubes were used per strain (6 replicates per timepoint). INH was added after 24, 48, 72, or 96 hours to a final concentration of 2.4 μg/mL. For each timepoint, CFUs were measured before adding INH (time 0 for each timepoint) and after 2 days of incubation with INH. The fraction of survival in INH was determined as the ratio between CFUs at day 2 over CFUs at time 0.

### RNA purification and quantitative PCR

For RNA purification from Mtb mc^2^6230, frozen cultures stored at −80°C were thawed on ice and centrifuged at 4,000 rpm for 5 min at 4°C. For Mtb H37Rv, cultures were pelleted and processed immediately. The pellets were resuspended in 1 mL Trizol (Life Technologies) and placed in tubes containing Lysing Matrix B (MP Bio). Cells were lysed by bead-beating (2 cycles of 9 m/sec for 40 s, with 2 min on ice in between) in a FastPrep 5G instrument (MP Bio). 300 μL chloroform was added and samples were centrifuged for 15 min at 4,000 rpm at 4°C. The aqueous phase was collected, and RNA was purified using Direct-Zol RNA miniprep kit (Zymo) according to the manufacturer’s instructions. For Mtb mc^2^6230 samples, the optional on-column DNase treatment step was used. For H37Rv samples, in-tube DNase treatment was done using DNase Turbo (Ambion) followed by purification with a Zymo Clean & Concentrator kit.

For cDNA synthesis, 600 ng of RNA were mixed with 0.83 μL 100 mM Tris, pH 7.5, and 0.17 μL of random primers (3 mg/mL NEB) in a total volume of 5.25 μL. The mix was denatured at 70°C for 10 min and placed on ice for 5 min. For reverse transcription, the following reagents were added to achieve the specified amounts or concentrations in 10 µL reactions: 100 U ProtoScript II reverse transcriptase (NEB), 10 U RNase inhibitor (murine; NEB), 0.5 mM each deoxynucleoside triphosphate (dNTP), and 5 mM dithiothreitol (DTT). Samples were incubated for 10 min at 25 °C and reverse transcription was performed overnight at 42°C. RNA was degraded by addition of 10 μL containing 250 mM EDTA and 0.5 N NaOH and heating at 65°C for 15 min, followed by addition of 12.5 μL of 1 M Tris-HCl, pH 7.5. cDNA was purified using the MinElute PCR purification kit (Qiagen) according to the manufacturer’s instructions.

RNA abundance was determined by quantitative PCR (qPCR) using iTaq SYBR green (Bio-Rad) with 200 pg of cDNA and 0.25 μM each primer in 10 μL reaction mixtures, with 40 cycles of 15 s at 95°C and 1 min at 61°C (Applied Biosystems 7500). Primers used in this study are listed in Table S2.

### Measurement of RNA half-life

For Mtb mc^2^6230, 5 mL of biological triplicates of log phase cultures were treated with RIF (final concentration of 50 μg/mL) to inhibit transcription, as reported by (Rustad *et al*., 2013). After RIF addition, tubes were placed into liquid nitrogen after 0, 2, 5, 10, 20, 40, 80, and 160 min. Cultures were frozen at −80 °C until RNA purification. RNA was purified using Direct-zol kit (Zymo) as described above. Half-lives were calculated as previously described (Vargas-Blanco *et al*., 2019, Nguyen *et al*., 2020). Timepoint 160 min was excluded from half-life calculations as it did not follow the initial exponential decay trend.

### Construction and analysis of RNA expression and 5’ end-directed libraries

For RNAseq libraries, 5 mL of log phase cultures of Mtb H37Rv WT, Δ*rnj*, and Δ*rnj::rnj*OE strains grown in 7H9 were placed in liquid nitrogen and stored at −80 °C. RNA purification and construction of both RNA expression and 5’ end-directed (non-5’ pyrophosphohydrolase-converted) libraries, were performed as previously reported (Martini *et al*., 2019, Martini *et al*., 2021). For both types of libraries, Illumina HiSeq 2000 paired-end sequencing producing 50 nt reads was used. Sequencing was performed at the UMass Medical School Deep Sequencing Core Facility. Raw and processed data are available in GEO, accession number GSE196357.

### Bioinformatic tools and analyses

Reads from Mtb datasets were aligned to the NC_000962 reference genome using Burrows-Wheeler Aligner (Li & Durbin, 2009). The FeatureCounts tool was used to assign mapped reads to genomic features, and DESeq2 was used to assess changes in gene expression in RNA expression libraries (Liao *et al*., 2014, Love *et al*., 2014). To differentiate fully upregulated genes from those having accumulation of reads in specific parts of the transcripts, we designed a pipeline (Figure S1). Each gene was divided into non-overlapping 10-nucleotide (nt) windows and the mean read depth (coverage) in each window was calculated for Δ*rnj* and WT replicates, using the Bedtools Genomecov function with the -pc option to computationally fill in coverage between reads (Quinlan & Hall, 2010), and a custom Python script to compute the coverage for each 10 nt window. The log_2_ ratio of coverage in Δ*rnj* /WT was computed for each window in each gene. Then, within each gene, the window with the median log_2_ coverage was identified. DESeq2 also reports log_2_ ratios for each gene, and the genome-wide mean of these log_2_ ratios was close to zero as expected given the assumption that the majority of genes do not differ between strains. However, the genome-wide mean of the median 10 nt window log_2_ ratios was −0.07. We therefore normalized the log_2_ ratios of the median 10 nt windows for all genes by adding 0.07. Next, we determined the absolute difference between the DESeq2 log_2_ fold change and the normalized median 10 nt window log_2_ ratio for each gene. The standard deviation (SD) of these differences was then calculated. Among the genes reported as differentially expressed by DESeq2, we set a cutoff of two SDs, such that genes with differences of ≥2 SDs were classified as partially up or down-regulated, and genes with differences of < 2 SD were classified as fully up or down-regulated.

To evaluate the characteristics of RNA fragments that accumulated in the Δ*rnj* strains, the normalized read depths for 5’ ends in the 5’ end-directed non-pyrophosphohydrolase-treated libraries were compared. 5’ ends were considered if they had a minimum read depth of five reads and the highest read depth in a 5-nt window (Table S5). Those 5’ ends with read depth ratios ≥10 in Δ*rnj* /WT were classified as enriched in the Δ*rnj* strain. A total of 381 5’ ends were classified as enriched. For a comparison, we selected 1,000 5’ ends that were equally represented in the WT and Δ*rnj* strains. The first 50 nt downstream of each 5’ end were analyzed. The minimum free energy (MFE) of each 50-nt sequence was determined using the RNAfold web server (Gruber *et al*., 2008), and the %GC was determined.

## RESULTS

### Mutations in RNase J are associated with drug resistance in clinical Mtb strains

We recently identified mutations in non-canonical drug resistance genes that are strongly associated with drug resistance in clinical Mtb strains isolated from China (Hicks *et al*., 2018). As mutations in these genes do not confer high-level drug resistance, we hypothesized that they may arise as intermediate steps in the acquisition of high-level drug resistance. One of the genes associated with drug resistance was Rv2752c, hereafter referred to as *rnj*, which encodes the bifunctional exo/endoribonuclease RNase J (Taverniti *et al*., 2011). Genome-wide association studies (GWAS) revealed that mutations in *rnj* were associated with INH resistance in Mtb (Hicks *et al*., 2018). Our results were consistent with two previous GWAS studies that also identified *rnj* mutations in association with multidrug resistance in Mtb (Zhang *et al*., 2013, Farhat *et al*., 2019).

In our data set, *rnj* was highly polymorphic in INH-resistant strains compared to drug sensitive strains and mutations were distributed throughout the CDS (Figure 1A, Table S3). Nonsynonymous mutations were the most prevalent (∼80% of the total mutations), while frameshift, and nonsense mutations were also found throughout the gene, suggesting selection for loss of function variants. Bellerose and collaborators (Bellerose *et al*., 2019, Bellerose *et al*., 2020) recently found, using a TnSeq screening approach, that disruption of *rnj* increased survival of Mtb in mice treated with the clinical first line drug regimen. We therefore hypothesized that loss of RNase J function might confer a survival advantage to the bacterium in the presence of drug treatment, and that mutations in clinical strains arise as a part of an evolutionary pathway to the acquisition of high-level drug resistance.

**Figure 1.**
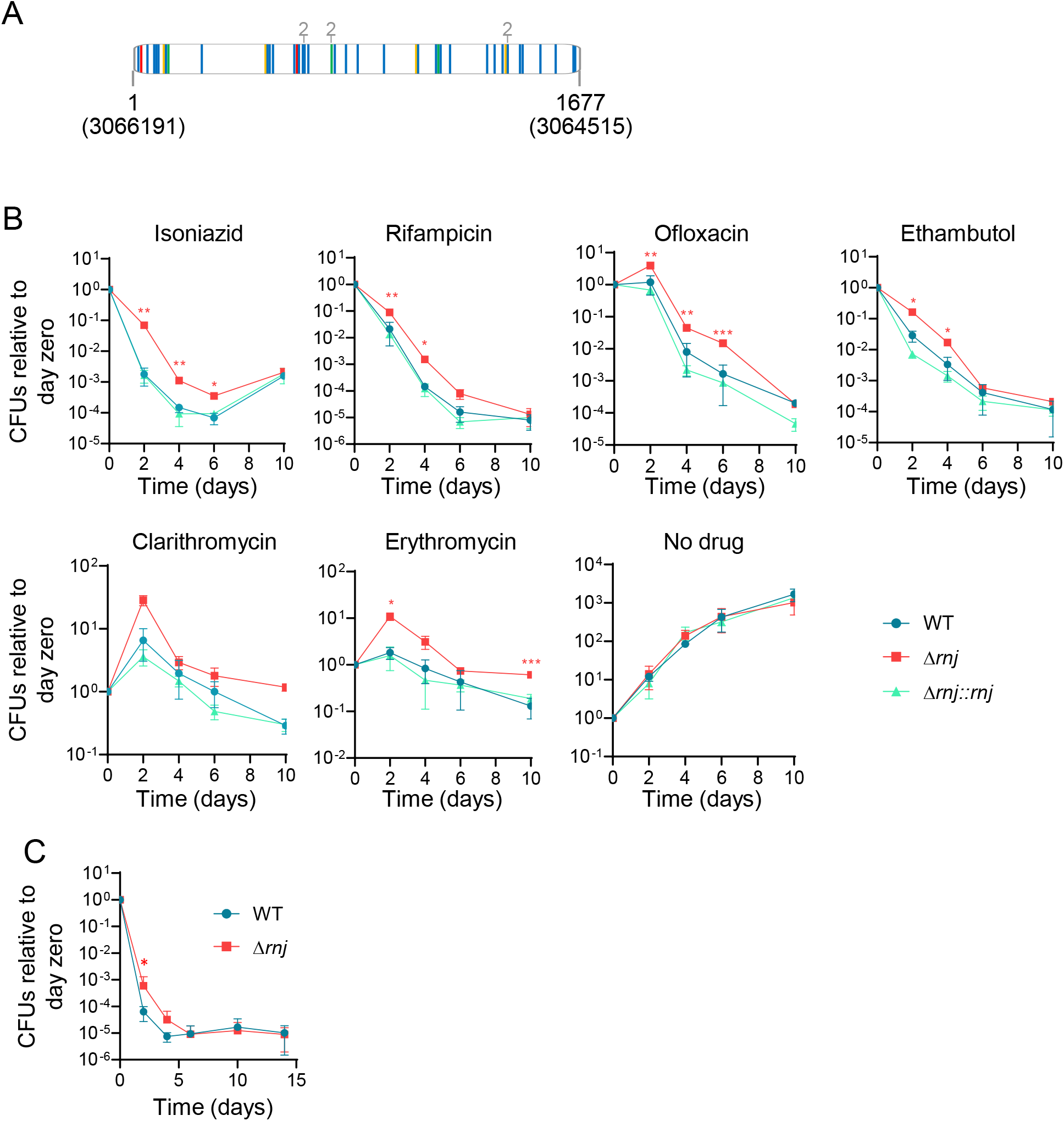
RNase J is mutated in many clinical Mtb strains, and loss of RNase J increases tolerance to several drugs. **A.** Vertical lines indicate mutations identified in clinical isolates in Hicks et al, 2018. Frameshift (red), nonsynonymous (blue), synonymous (orange), and nonsense (green) mutations are highlighted. Positions in the Mtb H37Rv genome are indicated in parenthesis. Numbers in grey indicate mutations that evolved twice independently. **B.** Time-kill curves in presence of the indicated drugs. The concentrations used were: 0.6 μg/mL RIF, 2.4 μg/mL INH, 2.5 μg/mL CLA, 1 μg/mL OFX, 2 μg/mL EMB, or 500 μg/mL ERY. **C.** Time-kill curves in presence of lethal concentrations of both INH (2.4 µg/mL) and OFX (2 µg/mL). Both experiments were performed using Mtb mc^2^6230 strains. \**p*<0.05, \*\**p*<0.01, \*\*\**p*<0.001 two-way ANOVA for comparisons of Δ*rnj* to WT. FDR 0.05 (Benjamini and Hochberg). Curves are representative of at least two independent experiments.

### Loss of RNase J increases tolerance to several drugs

To investigate the link between RNase J and drug sensitivity in Mtb, we first constructed RNase J deletion (Δ*rnj*6230) and complemented strains (Δ*rnj*6230*::rnj*) in the Mtb mc^2^6230 background (*ΔpanCD*, ΔRD1) (Sambandamurthy *et al*., 2006). To determine the effects of RNase J on drug sensitivity, we assessed both MICs and bactericidal activities of clinically relevant drugs. The MICs for RIF and INH were the same for the Δ*rnj*6230 strain and its WT parent (0.125 µg/mL for both drugs), indicating that loss of RNase J does not confer resistance to either drug. However, the Δ*rnj*6230 strain displayed increased survival in the presence of lethal concentrations of RIF and INH compared to the WT and the complemented strains (Fig 1B), suggesting that the absence of RNase J leads to increased drug tolerance. We observed a similar behavior when exposing the strains to lethal concentrations of EMB, OFX, CLA, and ERY (Fig 1B).

We sought to verify these results in virulent H37Rv, constructing RNase J deletion (Δ*rnj*H37Rv) and complemented strains (Δ*rnj*H37Rv*::rnj*). We tested killing by RIF, INH, and OFX, and found that in each case, the Δ*rnj*H37Rv strain had a survival advantage compared to the WT and complemented strains (Fig. S2). Furthermore, complementation of Δ*rnj*H37Rv with *rnj*^H86A^, which is predicted to be catalytically dead (Li de la Sierra-Gallay *et al*., 2008), did not restore the WT sensitivity phenotypes for RIF, INH, or OFX, suggesting that loss of RNase J catalytic activity is likely responsible for the greater survival of the Δ*rnj*H37Rv strain (Fig S2).

To further characterize the survival advantage conferred by loss of *rnj*, we exposed cultures to a combination of drugs (INH and OFX) at lethal concentrations to prevent outgrowth of resistant mutants. We observed that both strains reached a plateau at a similar number of CFUs (Fig 1C), suggesting that loss of RNase J does not affect persistence but rather increases drug tolerance by reducing the killing rate.

We noted that while the growth rate of Δ*rnj*6230 in mid-log phase was statistically indistinguishable from that of the WT and complemented strains, Δ*rnj*6230 had a slightly longer lag phase than WT cells (Fig 2A). The lag phase delay was condition-dependent, as we did not observe it when growth curves were started at a higher OD (Fig S3A), or when the strains were grown in minimal media (Fig S3B). Consistent with the lag phase defect in 7H9 broth, we found that Δ*rnj*6230 colonies took longer to arise on 7H10 plates and were smaller than those of the parental strain (Fig 2B and 2C). To test the possibility that the better survival of Δ*rnj* was due to the subtle growth defect observed at low ODs, we sub-cultured Δ*rnj*6230 and WT cultures to a low OD, added INH at a lethal concentration after one, two, three, or four days, and compared the fraction of survival after two days of incubation with drug (Fig 2D). We observed that the higher fraction of survival for Δ*rnj*6230 was consistent regardless of the growth phase at which drug was added (Fig 2E). In addition, we performed RIF and INH time-killing curves in minimal media, where Δ*rnj*6230 does not show a longer lag phase, and differences in drug killing between WT and Δ*rnj*6230 were similar to those observed in 7H9 media (Fig S4).

**Figure 2.**
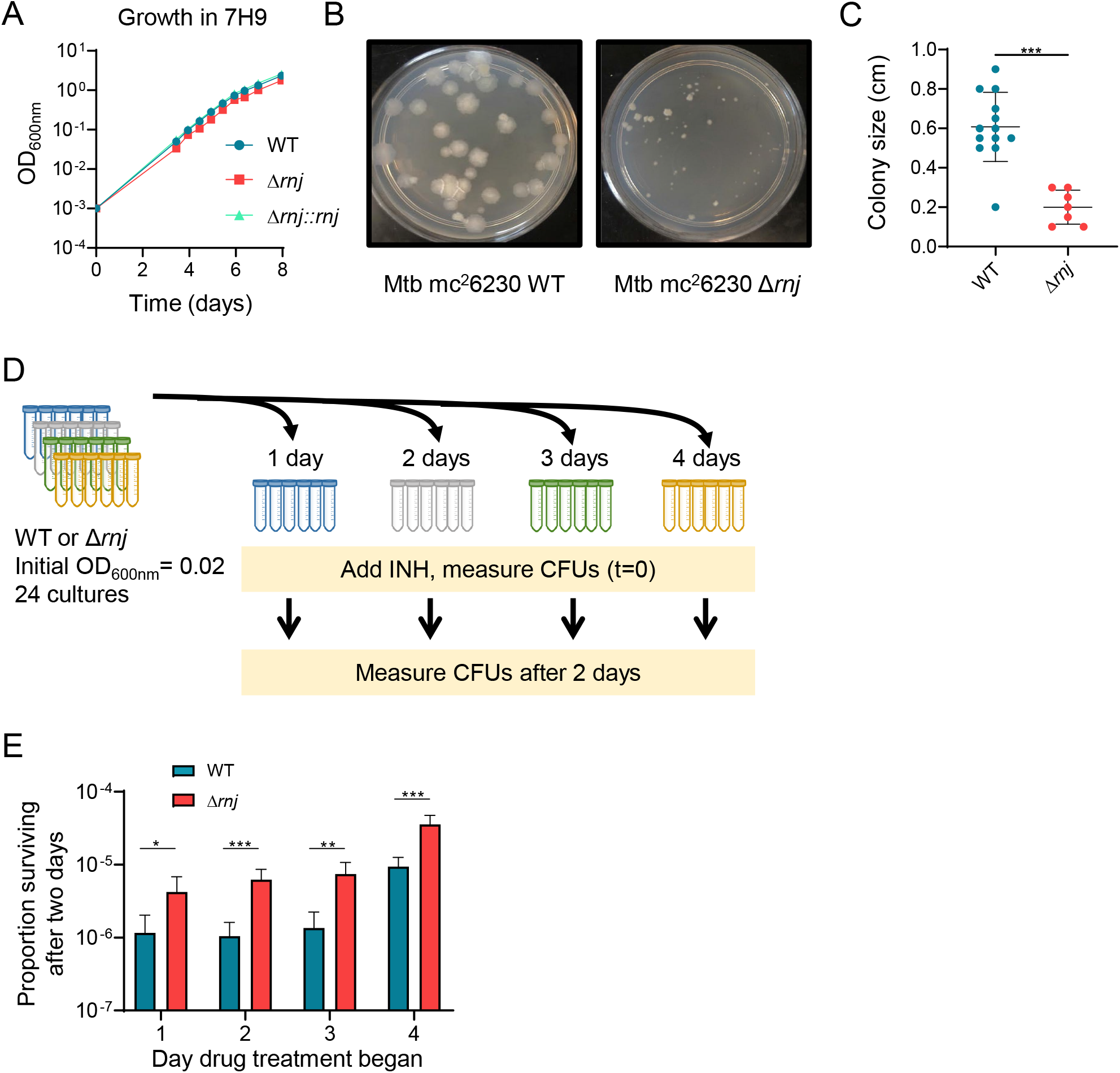
Drug tolerance in *Δrnj* Mtb is not due to its lag phase growth defect. **A.** Growth kinetics of Mtb mc^2^6230 WT, *Δrnj,* and Δ*rnj::rnj* in 7H9 media. Slopes are statistically equivalent for all three strains in mid-log phase (days 3-6, linear regression). ODs are significantly different for *Δrnj* vs WT for all time-points from day 3 onward (t-tests with FDR 1% correction) **B.** Colonies on 7H10 media 21 days after plating. **C.** Colony size was compared after 28 days of growth on 7H10 plates. \*\*\**p*<0.001, Mann-Whitney test. **D.** Schematic of experiment to determine the effect of growth phase on drug survival. **E.** Fraction of survival in INH (2.4 µg/mL) after two days of incubation with the drug starting at the indicated days. \**p*<0.05, \*\**p*<0.01, \*\*\**p*<0.001.

### RNase J affects rRNA processing and expression of a subset of genes and mRNA fragments

To investigate how RNase J mediates drug tolerance in Mtb, we assessed its role in RNA metabolism by performing RNAseq expression profiling as well as RNA 5’ end mapping. These studies were done with WTH37Rv and Δ*rnj*H37Rv strains transformed with the empty vector pJEB402 as well as the Δ*rnj*H37Rv strain complemented with *rnj* under a strong constitutive promoter, Δ*rnj*H37Rv*::rnj*OE (Ehrt *et al*., 2005). As a quality control, we evaluated the 23S rRNA as previous work has shown that RNase J is necessary for 23S rRNA maturation in *M. smegmatis* (Taverniti *et al*., 2011). Consistent with these data, we found that the 23S rRNA transcript was 15 nt longer in Δ*rnj*H37Rv compared to WTH37Rv (Fig S5), indicating that RNase J also plays a role in 23S rRNA processing in Mtb.

Standard analysis of the RNAseq expression data using DESeq2 indicated that 57 and 16 genes had increased or decreased transcript abundance, respectively, in Δ*rnj*H37Rv compared to WTH37Rv (fold change ≥1.5, adj *p* value ≤0.01) (Figure 3A, Table S4). Comparison of changes in transcript abundance in the Δ*rnj*H37Rv and Δ*rnj*H37Rv*::rnj*OE strains showed a significant negative correlation (Fig S6), reflecting the opposite effects of *rnj* overexpression and deletion. However, visual inspection of RNAseq expression library coverage revealed that several differentially abundant transcripts did not have increased abundance across the entire gene in Δ*rnj*H37Rv, but rather displayed increased read coverage for only short segments of the genes in question (Fig 3B). We hypothesized that these short RNA fragments might have arisen from incomplete degradation of mRNAs in the absence of RNase J. Thus, standard DESeq2 analysis is not sufficient to discriminate between genes for which the abundance of the whole transcript changes and genes marked by the accumulation of RNA fragments. To address this issue, we developed a bioinformatics pipeline (see Materials and Methods and Fig S1) to identify genes for which the abundance of the whole transcript was altered. Of the 57 genes that were reported as significantly overexpressed in Δ*rnj*H37Rv by DESeq2, 31 reflected increases in the whole transcript (Fig 3A, red dots and Table S4) while the remaining genes had increased abundance of only subsections of their transcripts (Fig 3A, light pink dots, and Table S4). All the genes with reduced abundance in Δ*rnj*H37Rv except for one (Fig 3A, light blue dot) showed changes in the abundance of the entire transcript (Fig 3A, blue dots, and Table S4).

**Figure 3.**
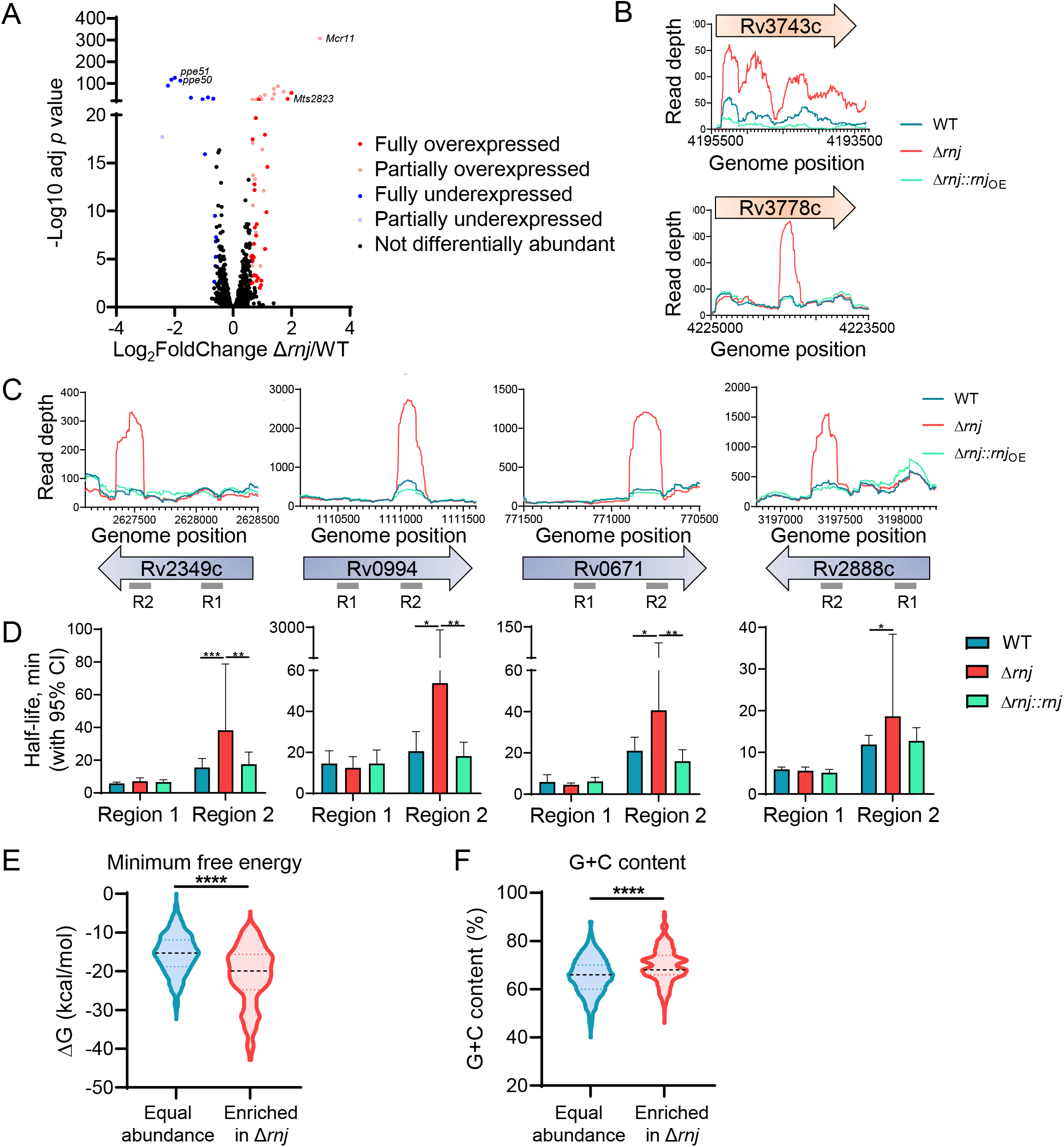
RNase J affects expression of genes and causes highly structured mRNA fragments to accumulate in Mtb. **A.** Volcano plot showing the genes affected by RNase J in Mtb H37Rv strains. Partially and fully over/under expressed genes are distinguished with different colors. **B.** Schematics of the read depth of two genes presenting full overexpression (upper panel) or partial overexpression (lower panel) in Δ*rnj*. **C.** Read depth of expression libraries in Mtb H37Rv strains for four genes that displayed partial overexpression in Δ*rnj*. Grey lines below the arrows denote the sequences targeted by qPCR for regions of the genes displaying accumulation of short fragments in Δ*rnj* (R2) and regions with similar read coverage in all strains (R1). For all H37Rv experiments, the WT and Δ*rnj* strains contained the empty vector pJEB402. **D.** Determination of half-life for the gene regions shown in C using Mtb mc^2^6230 strains. \**p*<0.05, \*\**p*<0.01, \*\*\**p*<0.001. **E** and **F.** 5’ end-mapping libraries were used to identify transcripts overexpressed in Δ*rnj*. The 50 nt downstream of both overexpressed and unchanged 5’ ends were analyzed to predict the minimum free energy of secondary structure formation and determine G+C content. \*\*\*\**p*<0.0001, Kruskal-Wallis test.

### RNase J has a specialized role in degrading mRNA fragments with strong secondary structure and high G+C content

We hypothesized that the transcript fragments that accumulated in the Δ*rnj*H37Rv strain could be RNase J targets that were inefficiently degraded in the mutant strain. To test this, we chose four genes with partial-transcript abundance increases in the Δ*rnj*H37Rv strain and measured the half-lives of both the overrepresented regions of each transcript and regions for which read coverage was similar in both strains (Fig 3C). The overrepresented regions had longer half-lives in the absence of *rnj*, while half-lives of equally abundant regions did not differ between the strains (Fig 3D).

To understand why RNase J has an apparently rate-limiting role in degradation of certain mRNA fragments, we assessed the GC content and predicted secondary structure of the 50 nt sequences adjacent to RNA 5’ ends that had increased abundance in the Δ*rnj*H37Rv strain or were equally abundant in the Δ*rnj*H37Rv and WTH37Rv strains. We found that the 5’ end-adjacent sequences with increased abundance in Δ*rnj*H37Rv had significantly more negative predicted minimum free energies of folding (ΔG) than those that were equally represented in Δ*rnj*H37Rv and WTH37Rv (Fig 3E). In addition, the median G+C content of the sequences overrepresented in Δ*rnj*H37Rv was significantly higher than that of the equally represented sequences (Fig 3F). These results are consistent with the idea that RNase J targets RNAs with relatively strong secondary structure.

Finding that loss of RNase J increased stability of short RNA fragments, we wondered if the increased abundance of the fully overexpressed genes in Δ*rnj*H37Rv was a direct consequence of slower degradation rates or increased transcription. We measured the half-lives of six fully overexpressed genes and found that transcript stability was not altered (Fig S7). We also considered the possibility that the absence of RNase J could lead to accumulation of antisense transcripts that could affect mRNA degradation or translation. However, there were no changes in median antisense coverage for genes differentially expressed in Δ*rnj*H37Rv or for all expressed genes in this strain. Together, these data suggest that RNase J affects the transcript abundance of some genes through altered transcription rather than by altering their mRNA stability.

### Genes differentially expressed in the absence of RNase J are enriched for sRNAs, PE/PPE family genes, SigM targets, and genes with roles in hypoxia response and carbon source switching

Examination of the genes that were fully overexpressed or underexpressed revealed several themes. First, the differentially expressed genes were enriched for sRNAs and genes of the PE/PPE family, including nine overexpressed PE_PGRS genes (Fig S8). Second, six of the underexpressed genes (*ppe50*, *ppe51*, *fadD26*, *ppsA*, *Rv0885*, and *Rv3137*) were reported to be negatively regulated by the stress-responsive alternative sigma factor SigM, while one of the overexpressed genes (*Rv3093c*) was reported to be positively regulated by SigM (Raman *et al*., 2006). This suggests that there may be increased SigM activity in Δ*rnj*H37Rv, although the *sigM* gene itself was not increased at the transcript level. Finally, the differentially expressed genes included several associated with hypoxia responses (the sRNAs MTS2823 and F6, and the protein-coding genes *fdxA*, *ppe31*, *clgR,* and *Rv3740c*) and several associated with utilization of various carbon sources (*ppe50*, *ppe51, mcm1C*, *prpC*, *Rv1066*) (Park *et al*., 2003, Boshoff *et al*., 2004, Daniel *et al*., 2004, Muñoz-Elías *et al*., 2006, Forrellad *et al*., 2014, Rodríguez *et al*., 2014, Del Portillo *et al*., 2018, Korycka-Machała *et al*., 2020, Dechow *et al*., 2021), suggesting that the metabolic status of the Δ*rnj*H37Rv may differ from that of WTH37Rv.

### Overexpression of the sRNAs *Mts2823* and *Mcr11* is necessary but not sufficient for INH tolerance in Δ*rnj* Mtb

The sRNAs *Mts2823* and *Mcr11* were two of the most overexpressed genes in the Δ*rnj*H37Rv strain (Table S4). Since sRNAs have been implicated in adaptation to different stresses, we sought to investigate if their increased expression in Δ*rnj* contributed to drug tolerance. We therefore deleted each of the two sRNAs in both the WT6230 and Δ*rnj*6230 strains and performed time-killing curves in presence of RIF or INH. We found that deletion of either of these sRNAs in the Δ*rnj*6230 background decreased INH tolerance to levels near the WT6230 (Fig 6A-B and 6D-E). By contrast deletion of these sRNAs had no effect on INH tolerance in WT6230 background (Fig 6C and 6F). Thus, *Mts2823* and *Mcr11* are necessary for the INH tolerance conferred by loss of RNase J in the Δ*rnj*6230 strain. To determine whether either sRNA was sufficient for INH tolerance, we then constructed strains overexpressing either *Mts2823* (*Mts2823*OE) or *Mcr11* (*Mcr11*OE) in the WT6230 background. However, we found that both showed drug sensitivity levels comparable to that of WT6230 (Fig 6C and 6F), suggesting that overexpression of either *Mts2823* or *Mcr11* alone is not sufficient to increase drug tolerance. No consistent effects were observed for strains with deletions or overexpression of these sRNAs in RIF, suggesting that effectors acting downstream of RNase J deletion may act in part via drug specific mechanisms (Fig S9).

### Downregulation of *ppe50-ppe51* is necessary for the INH and RIF tolerance phenotypes of Δ*rnj* Mtb

Two of the most strongly downregulated genes in the Δ*rnj*H37RV strain were *ppe50* and *ppe51*, which are expressed in an operon and have previously been implicated in drug sensitivity (Xu *et al*., 2017, Bellerose *et al*., 2019, Bellerose *et al*., 2020). To test the hypothesis that downregulation of *ppe50* and *ppe51* contributes to the drug tolerance of the Δ*rnj* strains, we ectopically overexpressed them in the Δ*rnj*6230 background and performed drug killing experiments. Overexpression of the *ppe50-ppe51* operon in the Δ*rnj*6230 strain impaired survival of Mtb in the face of both INH and RIF, producing drug sensitivity levels similar to those observed in the WT strain (Fig 5). Thus, the dysregulated expression of *ppe50-ppe51* is also necessary for the altered drug susceptibility associated with loss of *rnj*.

**Figure 4.**
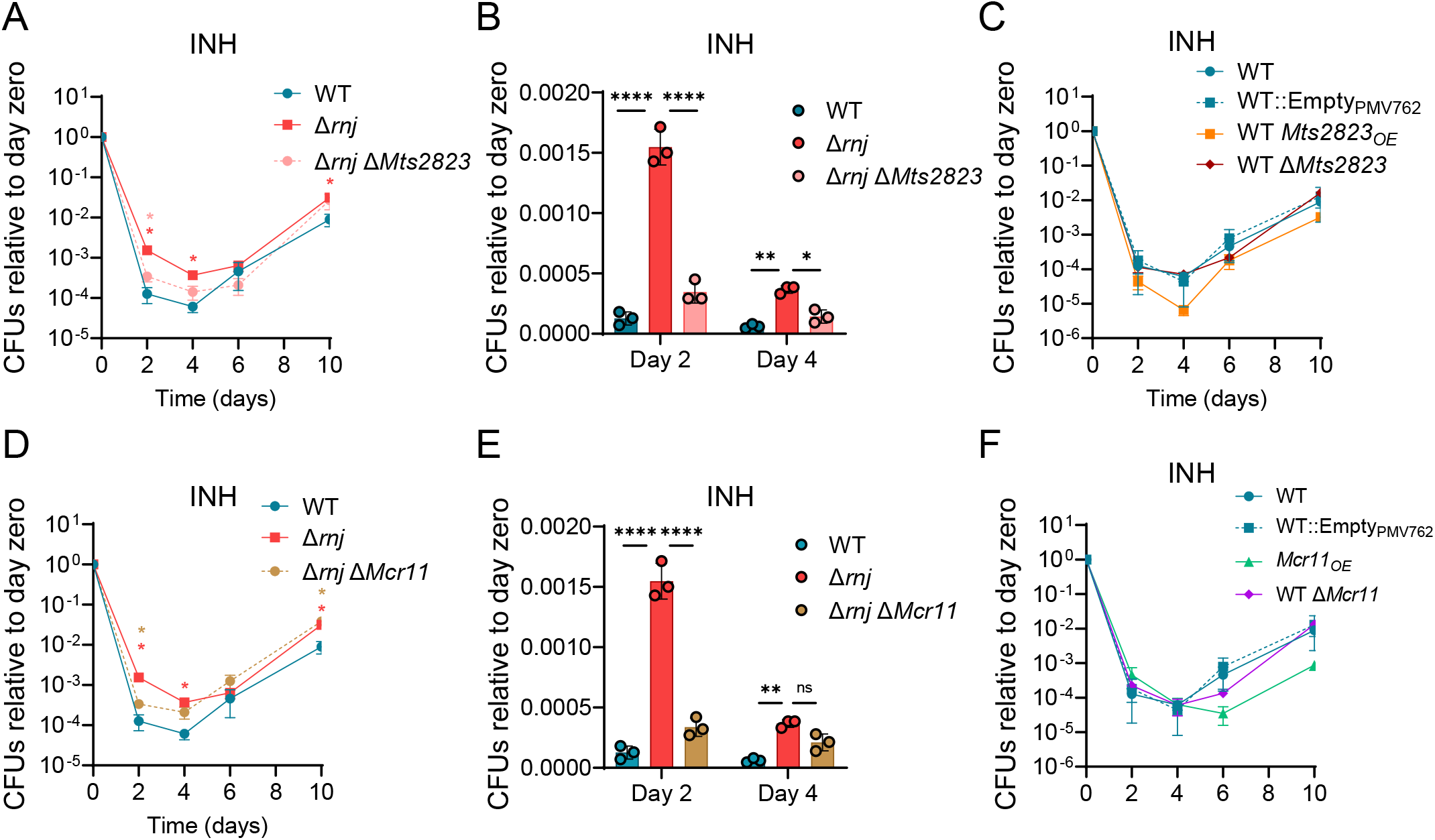
Overexpression of the sRNAs *Mts2823* and *Mcr11* is necessary but not sufficient for INH tolerance in Δ*rnj* Mtb. Time-killing curves in presence of INH (2.4 µg/mL) for strains with deletion or overexpression of *Mts2823* **(A-C)** or *Mcr11* **(D-F)** strains are shown. \**p*<0.05, \*\**p*<0.001 two-way ANOVA. Pink stars: comparison of Δ*rnj* Δ*Mts2823* to Δ*rnj*. Red stars: comparison of WT to Δ*rnj.* Tan stars: comparison of Δ*rnj* Δ*Mcr11* to Δ*rnj*.

**Figure 5.**
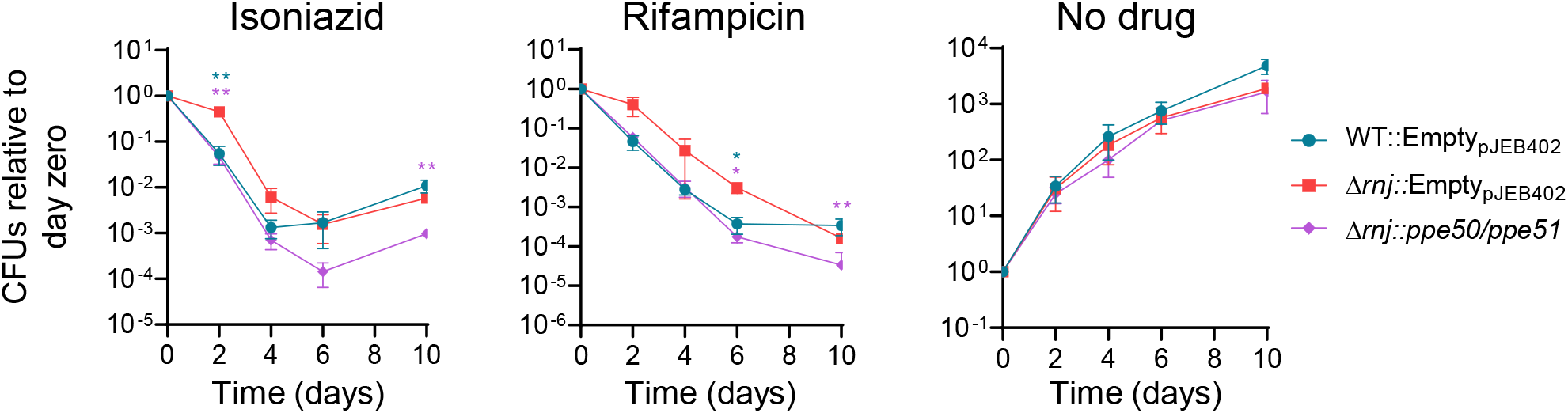
Downregulation of *ppe50-ppe51* is required for the drug tolerance phenotype of Δ*rnj* Mtb. Time-kill curves in the presence of RIF (0.6 µg/mL) or INH (2.4 µg/mL) in Mtb mc^2^6230 strains. \**p*<0.05, \*\**p*<0.01, two-way ANOVA. Blue stars: comparison of WT::Empty_pJEB402_ to Δ*rnj::*Empty_pJEB402_. Violet stars: comparison of Δ*rnj::ppe50/ppe51* to Δ*rnj::*Empty_pJEB402_. FDR 0.05 (Benjamini and Hochberg).

## DISCUSSION

Previous studies have shown that mutations in the RNase J-encoding gene were more prevalent in drug resistant Mtb clinical isolates compared to drug-sensitive isolates (Zhang *et al*., 2013, Hicks *et al*., 2018, Farhat *et al*., 2019). Here we show that loss of RNase J function increases drug tolerance, an advantageous trait to overcome drug stress during TB treatment that could promote further development of high-level drug resistance in clinical settings. We have furthermore defined a role for RNase J in mycobacterial mRNA metabolism as a specialized degradation factor.

Despite the non-essentiality of RNase J in mycobacteria, a recent study showed that RNase J is a major component of the degradosome in Mtb (Płociński *et al*., 2019). Previous work also implicated RNase J as having roles in rRNA processing in *M. smegmatis* (Taverniti *et al*., 2011). In *M. abscessus*, deletion of the *rnj* gene (MAB_3083c) resulted in changes in colony morphology, biofilm formation, and sliding motility (Liu *et al*., 2021). However, the role of this nuclease in mycobacterial mRNA metabolism remained elusive. Our finding that deletion of RNase J affects expression of a limited number of genes is consistent with the near-wildtype growth characteristics of the mutant. Taken together with the essentiality of RNase E in mycobacteria, our results point to RNase E as playing a more important role in bulk mRNA decay. This is consistent with work in two other species that encode both RNase E and RNase J, *Rhodobacter sphaeroides* and *Synechocystis* sp. PCC6803, where the respective roles of the two nucleases have been investigated (Rische-Grahl *et al*., 2014, Cavaiuolo *et al*., 2020). In contrast, RNase J plays a leading role in bulk mRNA degradation in several species that naturally lack RNase E, such as *B. subtills*, *C. diphtheriae, S. aureus* and *H. pylori* (Durand *et al*., 2012, Linder *et al*., 2014, Redko *et al*., 2016, Luong *et al*., 2021).

Our analyses allowed us to identify native RNase J targets in Mtb as having higher G+C content and stronger predicted secondary structure than non-targets. It is important to note that the mRNA fragments we identified as direct RNase J targets generally had longer half-lives than other fragments of the same transcript even when RNase J was present (Fig. 3D, comparison of region 1 to region 2 in WT strain), in agreement with the idea that RNase J targets RNAs that are refractory to degradation by the bulk degradation machinery. This role for mycobacterial RNase J is consistent with a recently described duplex-unwinding activity in the archaeal mpy-RNase J, which was shown to degrade highly structured RNAs *in vitro* (Li *et al*., 2020).

We demonstrated here that deletion of *rnj* reduces bacterial killing when Mtb is exposed to lethal concentrations of several drugs and proved that these observations are due to an increase in drug tolerance rather to a higher formation of persister cells. The drug tolerance is explained at least in part by gene expression changes. The overexpression of two PE/PPE genes that were downregulated in *Δrnj*, *ppe50-ppe51*, restored the RIF and INH sensitive phenotype in the RNase J mutant to the WT levels, demonstrating that downregulation of these genes is necessary for drug tolerance in *Δrnj*. With respect to INH, this finding is consistent with previous studies reporting that deletion of *ppe51* lead to increased bacterial survival in INH-treated mice (Bellerose *et al*., 2019, Bellerose *et al*., 2020) and reduced sensitivity to INH in vitro (Xu *et al*., 2017, Bellerose *et al*., 2020). The relationship between *ppe51* and RIF sensitivity appears to be more complex. Deletion of *ppe51* was previously found to cause a small but significant increase in the MIC for RIF *in vitro*, but led to increased RIF sensitivity in mice (Bellerose *et al*., 2020). The impact of *ppe51* on RIF sensitivity may therefore be condition-dependent.

It is possible that the effects of *ppe50-ppe51* on drug sensitivity are related to carbon metabolism. Dechow and collaborators reported that specific PPE51 variants promoted glycerol uptake and prevented growth arrest in acidic conditions when glycerol was the sole carbon source (Dechow *et al*., 2021). On the other hand, knockdown of *ppe51* affected uptake of disaccharides and attenuated Mtb growth in minimal media with disaccharides as the sole carbon source (Korycka-Machała *et al*., 2020). There is an increasing body of literature implicating glycerol metabolism as a pathway that affects sensitivity of Mtb to various drugs (Xu *et al*., 2017, Bellerose *et al*., 2019, Safi *et al*., 2019). It is therefore conceivable that reduced expression of *ppe51* in the *Δrnj* strains leads to INH tolerance through a mechanism related to glycerol metabolism.

Deletion of two non-coding sRNAs overexpressed in the mutant, *Mts2823* and *Mcr11*, in the *Δrnj* background also showed partial restoration of drug sensitivity, suggesting that RNase J likely modulates drug tolerance via multiple mechanisms. *Mts2823* is an sRNA orthologous to the *E. coli* 6S RNA, which interacts with the RNA polymerase core preventing gene expression in mycobacteria and is highly expressed in stationary phase (Arnvig *et al*., 2011, Hnilicová *et al*., 2014). Overexpression of *Mts2823* was shown to have a slight effect on the growth rate in Mtb (Arnvig *et al*., 2011), consistent with our observations for the *Mts2823*OE strain in absence of drug. *Mcr11* is highly expressed during mouse infection (Pelly *et al*., 2012). Interestingly, the expression of this sRNA was increased for only ∼80% of the transcript sequence, with the 3’ end region showing similar levels as in the WT strain (Fig S10), indicating that RNase J could be involved in maturation of the 3’ end of the transcript. However, more work needs to be done to understand the mechanisms of such regulation.

Taken together, our results suggest a scenario in which RNase J activity affects expression of multiple genes that together affect drug tolerance. Some of these expression changes have relatively well-delineated impacts (*e.g.*, loss or reduction of *ppe51* expression consistently causes reduced INH sensitivity in our work and that of others), while others appear to have different effects in WT and Δ*rnj* backgrounds. These findings, together with our observation that Δ*rnj* strains accumulate mRNA degradation intermediates, are consistent with the idea that mutations in *rnj* have pleiotropic effects on the physiology of Mtb and may therefore be selected *in vivo* in response to a variety of pressures.

## AUTHOR CONTRIBUTIONS

MCM, NDH, SSS, and SMF conceived and designed experiments. MCM, NDH, TB, and JS performed experiments. JX and SSS designed data analysis methods. JX and MCM performed data analysis. MCM, NDH, SSS, and SMF wrote the manuscript.

## ACKNOWLEDGEMENTS

We thank members of the Fortune and Shell labs and members of the Pathway Analysis in Tuberculosis P01 group for helpful feedback and discussions.

## DATA AVAILABILITY

Raw and processed RNAseq data are available in GEO, accession number GSE196357.

## Sup. Figures

**Figure S1.**
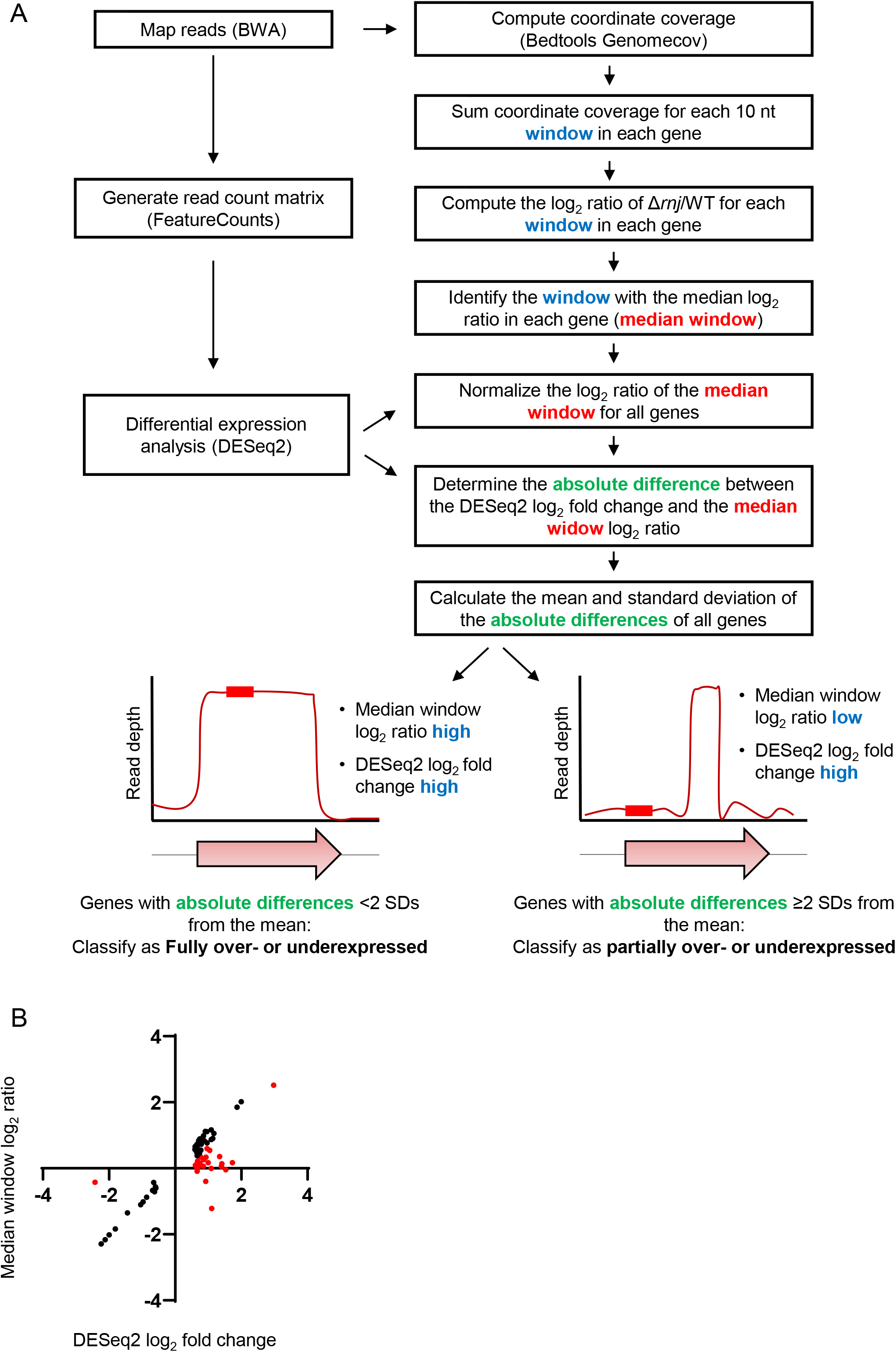
Bioinformatic analysis of RNASeq data. **A.** Schematic representation of the pipeline used to differentiate fully overexpressed genes from partially overexpressed genes. **B.** Scatter plot representing the relationship between DESeq2 log_2_ fold change and median 10 nt window log_2_ ratio. The red dots indicate partially under/over expressed genes and black dots indicate fully under/over expressed genes.

**Figure S2.**
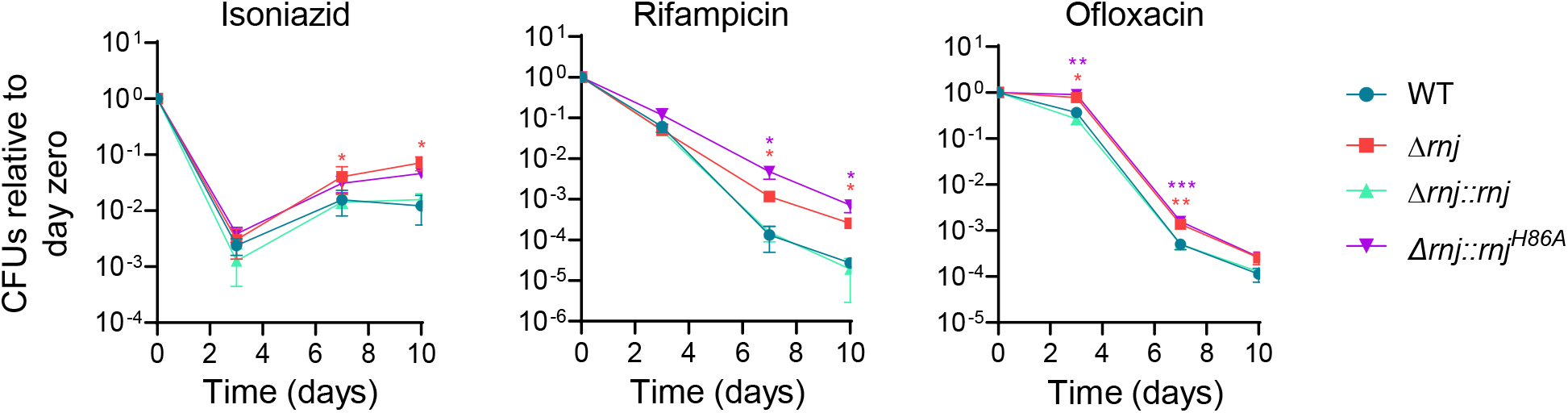
Loss of RNase J affects drug sensitivity in Mtb H37Rv. Time-kill curves comparing *Δrnj* transformed with empty vector (pJEB402), Δ*rnj* complemented with the catalytic site mutant *rnj^H86A^* under the strong UV15 promoter, Δ*rnj::rnj*, and the WT strain transformed with empty vector (pJEB402) in the H37Rv background are shown. \**p*<0.05, **p<0.01, \*\*\**p*<0.001 two-way ANOVA comparing Δ*rnj* (red stars), Δ*rnj::rnj* (green stars) and Δ*rnj::rnj^H86A^* (violet stars) to the WT (control group), with FDR 0.05 (Benjamini and Hochberg).

**Figure S3.**
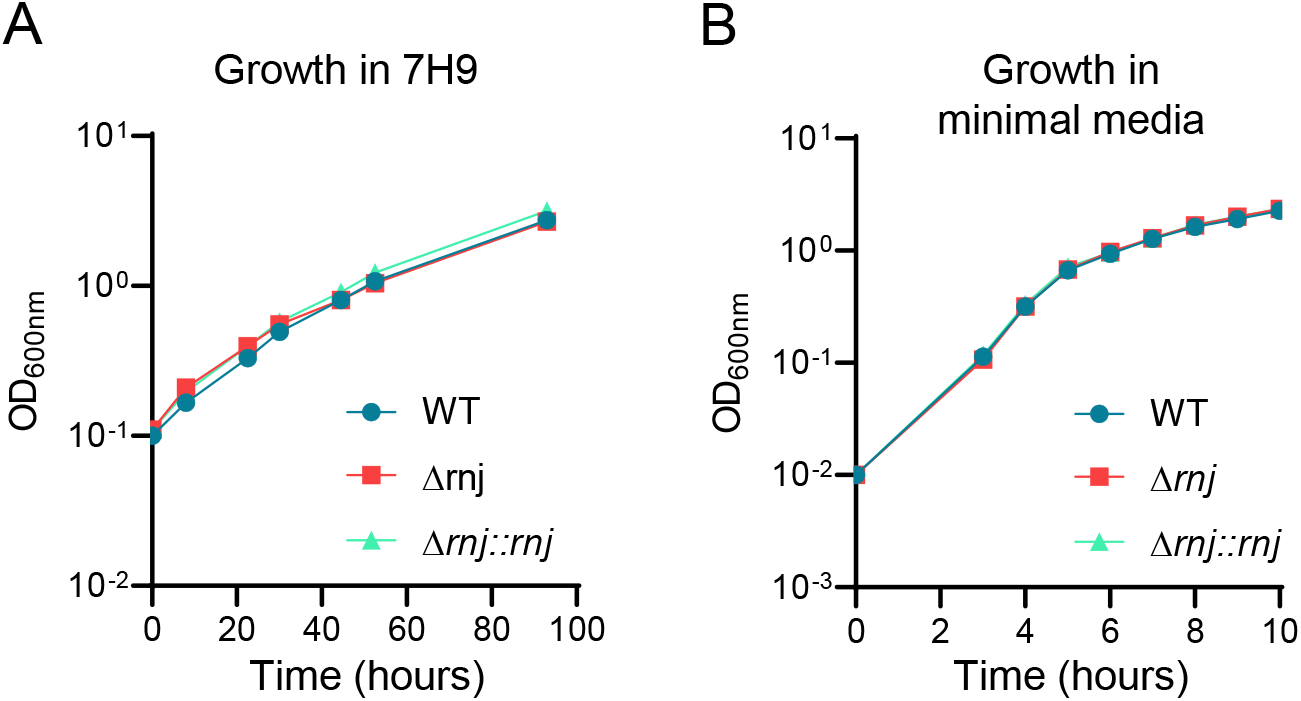
Growth kinetics in drug-free media. **A.** Mtb mc^2^6230 WT, *Δrnj,* and Δ*rnj::rnj* were grown in 7H9 media, starting with an initial OD_600nm_=0.1. **B.** Strains were grown in minimal media (initial OD_600nm_=0.01) supplemented with glycerol and tween-80.

**Figure S4.**
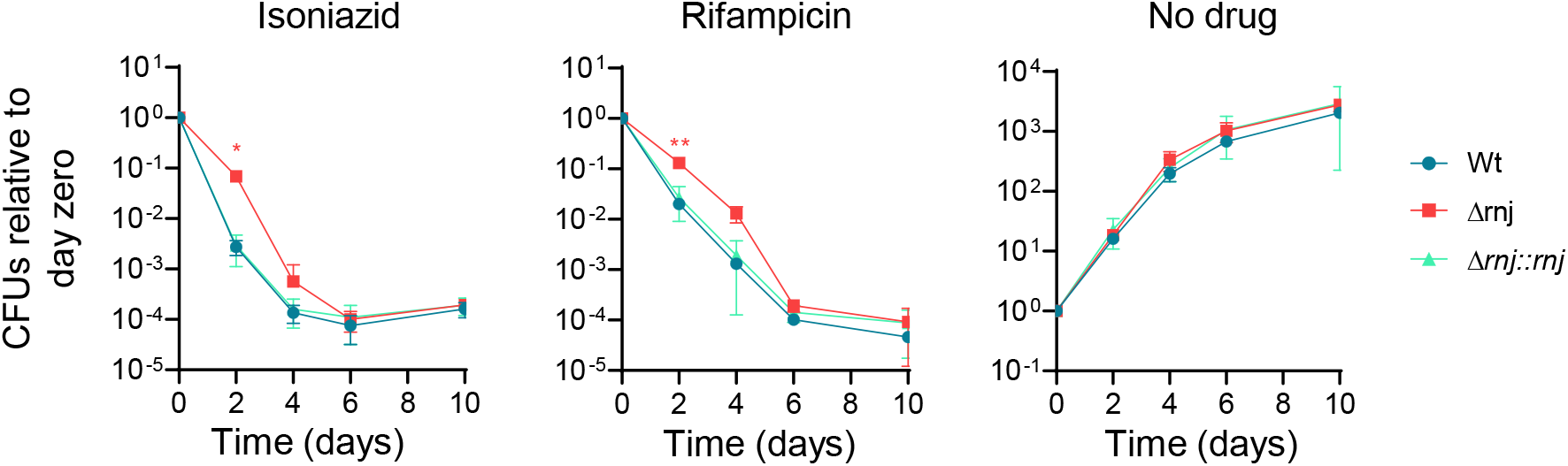
Time-kill curves in minimal media. Time-kill curves comparing *Δrnj* and Δ*rnj::rnj* to the WT strain in the Mtb mc^2^6230 background are shown. \**p*<0.05, \*\**p*<0.01 two-way ANOVA with FDR 0.05 (Benjamini and Hochberg).

**Figure S5.**
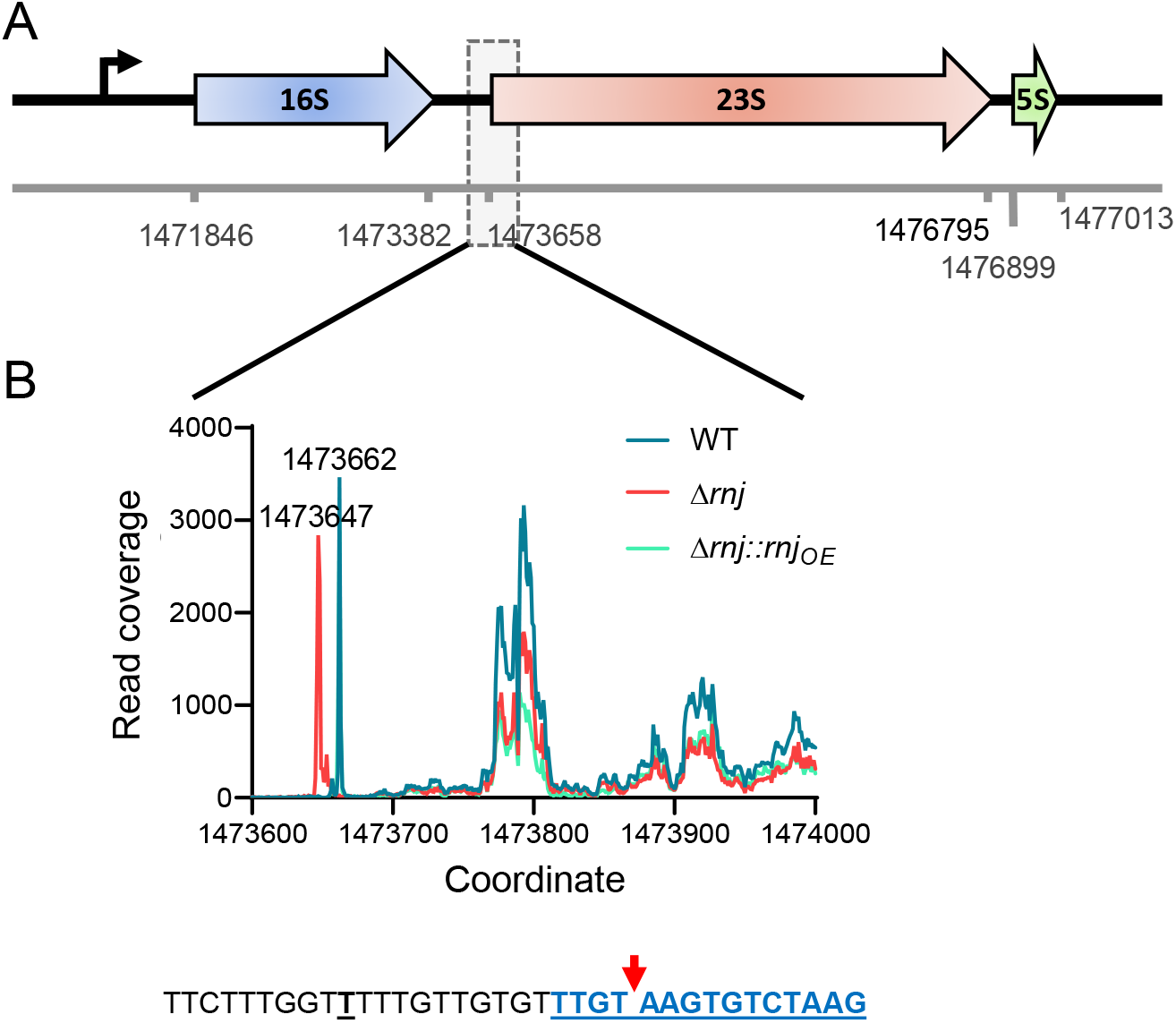
RNase J contributes to 23S rRNA maturation in Mtb. **A.** Schematic of the rRNA operon in Mtb. The genome positions in the reference genome NC_000962 are given. The RNase J-dependent processing region of the annotated 23S rRNA is shown in a dashed rectangle. **B.** Read depth of 5’ end-mapping libraries is shown for the indicated genome coordinates for Δ*rnj*, WT and Δ*rnj::rnj_OE_* strains. The WT and Δ*rnj* strains contained the empty vector pJEB402. The first part of 23S rRNA sequence is highlighted in underlined bolded blue. We did not detect the annotated 23S 5’ end at coordinate 1473658 (the first blue nt). The red arrow shows the identified processed 5’ end (genome coordinate 1473662) that predominated in the WT and complemented strains. The underlined black “T” indicates the 5’ end (genome coordinate 1473647) that predominated in the Δ*rnj* strain.

**Figure S6.**
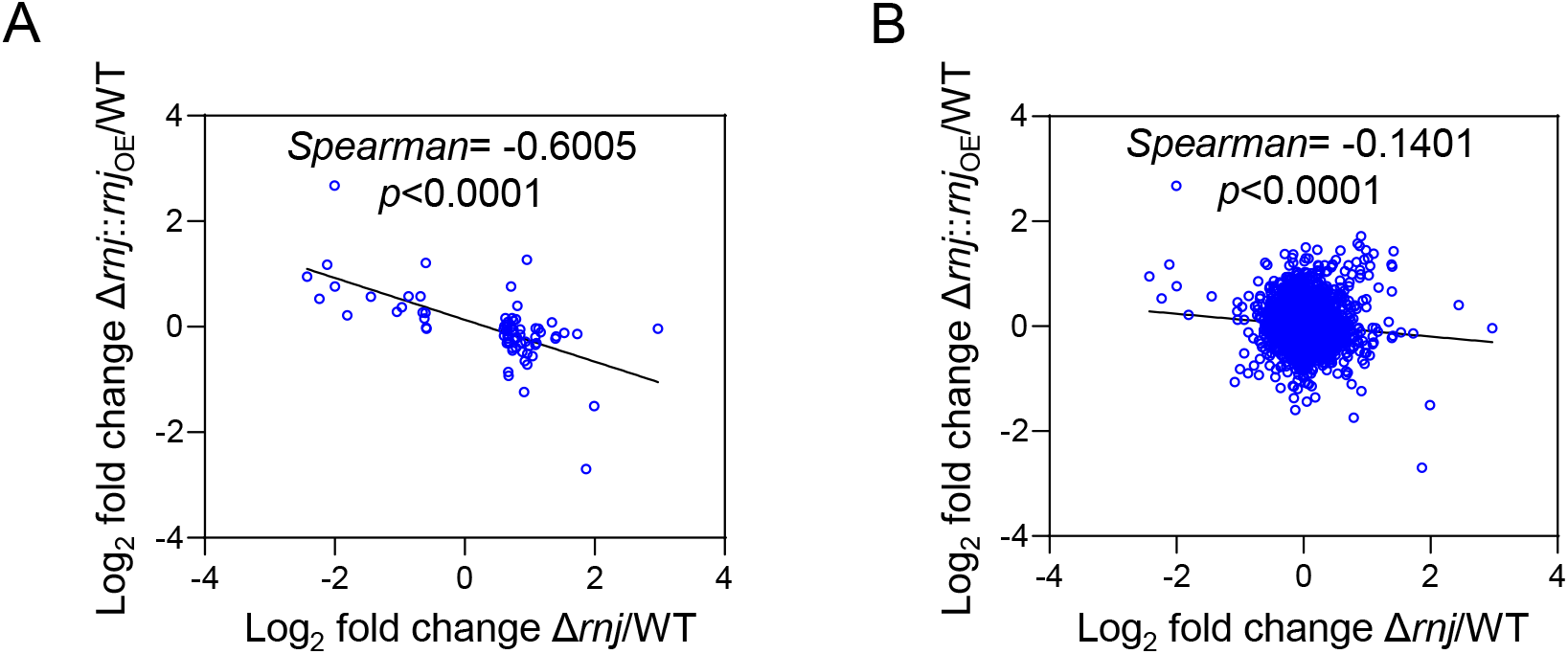
Genes affected by loss of *rnj* are inversely affected by *rnj* overexpression. DESeq2 analysis was performed independently for Δ*rnj*/WT and Δ*rnj::rnj*_OE_/WT and the outputs were compared. The WT and Δ*rnj* strains contained the empty vector pJEB402. In **A**, only those genes that were differentially expressed in Δ*rnj* are shown. In **B** all the genes used in DEseq2 analysis are shown.

**Figure S7.**
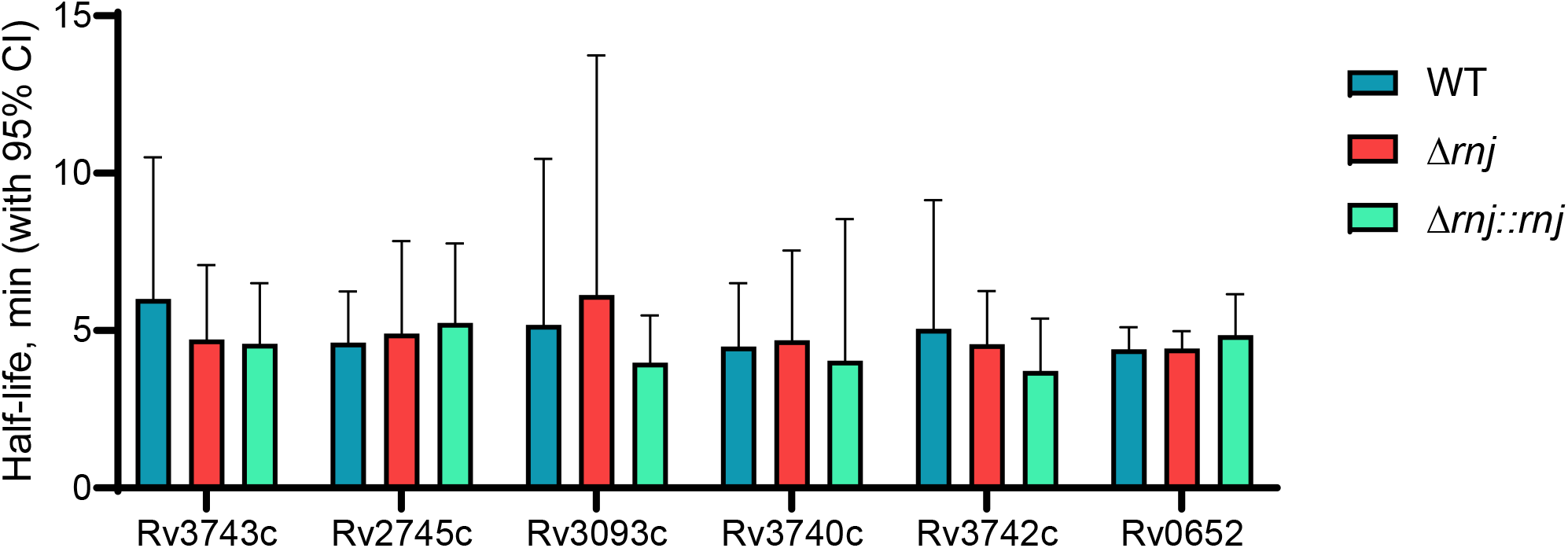
Some of the genes fully overexpressed in Δ*rnj* do not display increased stability. Half-lives of 6 genes fully overexpressed in Δ*rnj* were measured in Mtb mc^2^6230 WT, Δ*rnj,* and Δ*rnj::rnj* strains.

**Figure S8.**
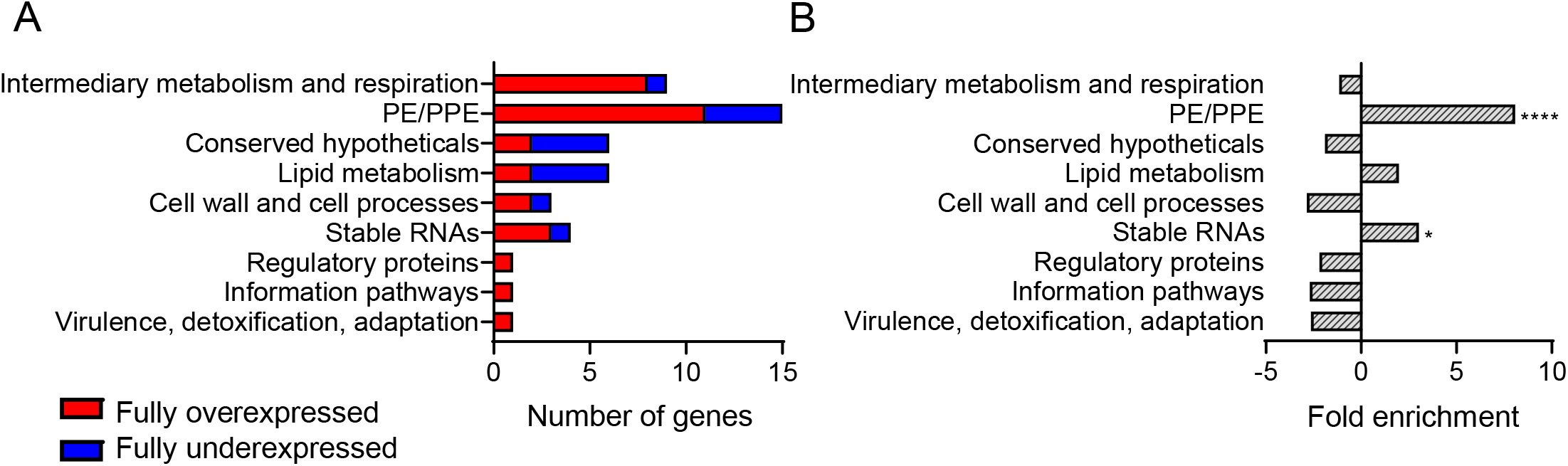
Genes that are differentially expressed in the absence of *rnj* in the H37Rv background are enriched for stable RNAs and PE/PPE family genes. A. Classification of the fully over- and underexpressed genes by category. **B.** Gene category enrichment for fully over- and underexpressed genes using hypergeometric test. \**p*<0.05, \*\*\*\**p*<0.001.

**Figure S9.**
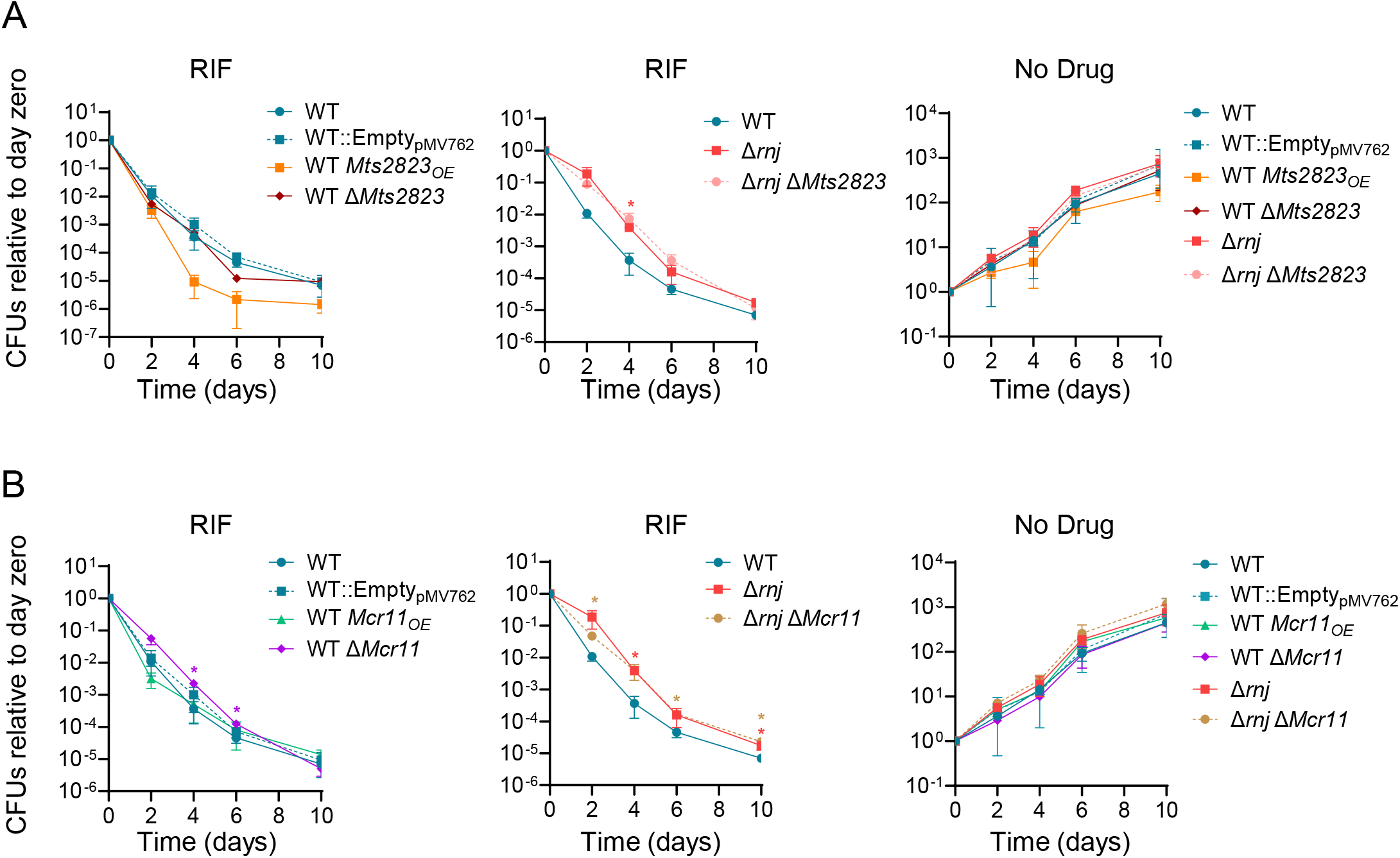
The sRNAs *Mts2823* and *Mcr11* do not affect RIF tolerance in WT or Δ*rnj* Mtb. Time-killing curves in presence RIF (0.6 µg/mL) for strains with deletion or overexpression of *Mts2823* **(A)** or *Mcr11* **(B)** strains are shown. OE indicates overexpression of the indicated sRNA. Growth in drug-free media was also assessed. \**p*<0.05, \*\**p*<0.001, two-way ANOVA. Red stars: comparison of WT to Δ*rnj*. Violet stars: comparison of WT Δ*Mcr11* to WT. Tan stars: comparison of Δ*rnj* Δ*Mcr11* to WT. The apparent increase in RIF tolerance seen here for the WT Δ*Mcr11* strain was not reproducible.

**Figure S10.**
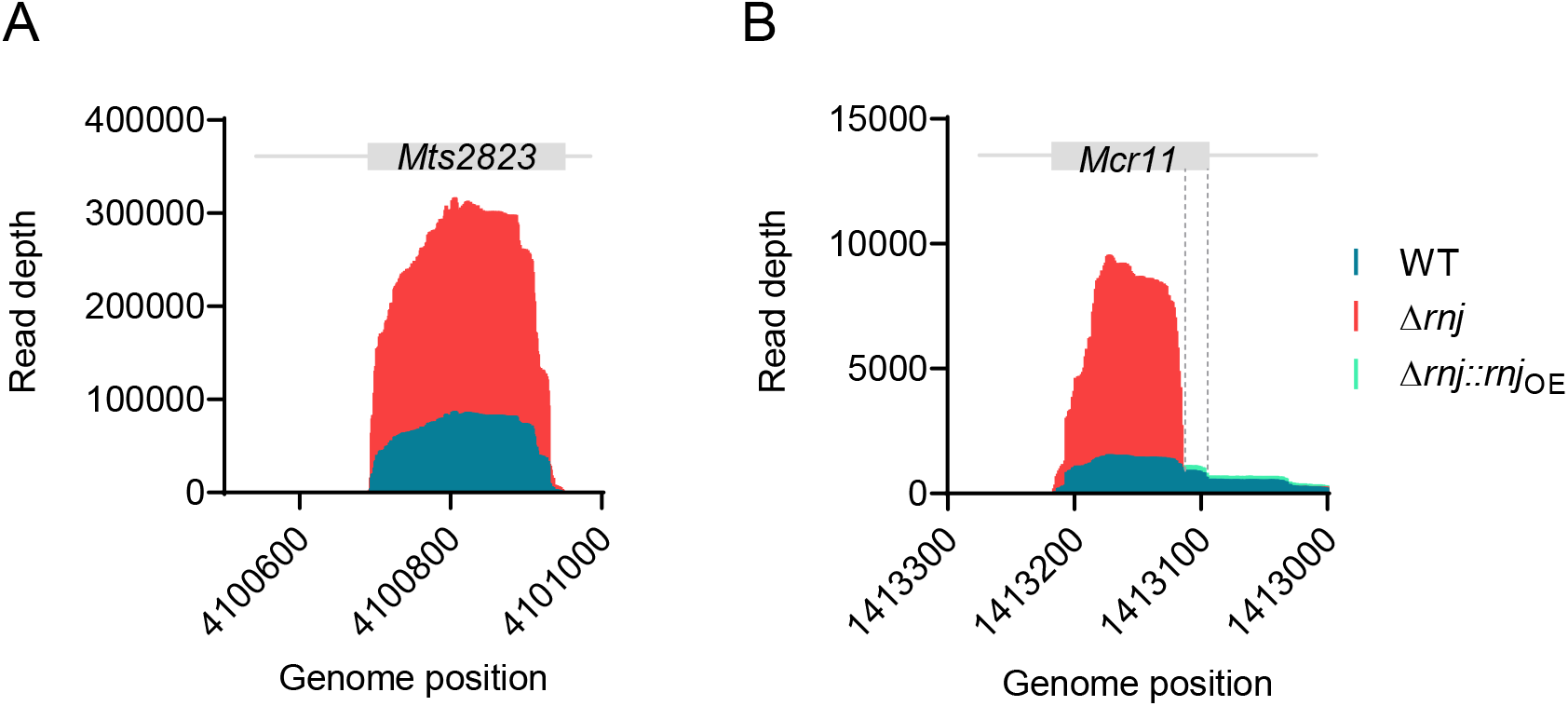
RNAseq coverage plots of sRNAs *Mts2823* (A) and *Mcr11* (B). Read depth in RNAseq expression libraries is shown. The region of *Mcr11* displaying similar coverage in Δ*rnj* and the WT is denoted with dashed lines. The WT and Δ*rnj* strains contained the empty vector pJEB402.

